# Temporal Dynamics of HCMV Gene Expression in Lytic and Latent Infections

**DOI:** 10.1101/2021.07.26.453763

**Authors:** Batsheva Rozman, Aharon Nachshon, Roi Levi Samia, Michael Lavi, Michal Schwartz, Noam Stern-Ginossar

## Abstract

Primary infection with Human cytomegalovirus (HCMV) results in a persistent lifelong infection due to its ability to establish latent infection. During productive HCMV infection, viral genes are expressed in a coordinated cascade that is characteristic of all herpesviruses and traditionally relies on the dependencies of viral genes on protein synthesis and viral DNA replication. In contrast, the transcriptional landscape associated with HCMV latency is still disputed and poorly understood. Here, we examine viral transcriptomic dynamics during the establishment of both productive and latent HCMV infections. Our temporal measurements reveal that viral gene expression dynamics along productive infection and their dependencies on protein synthesis and viral DNA replication, do not fully align. This illustrates that the regulation of herpesvirus genes does not represent a simple sequential transcriptional cascade, and surprisingly, many viral genes are regulated by multiple independent modules. Using our improved classification of viral gene expression kinetics in conjunction with transcriptome-wide measurements of the effects of a wide array of chromatin modifiers, we unbiasedly show that a defining characteristic of latent cells is the unique repression of immediate early (IE) genes. Altogether, our findings provide an elaborate definition of HCMV gene expression patterns and reveal novel principles that govern viral gene expression in lytic and latent infection states.

## Introduction

Human cytomegalovirus (HCMV) is a pervasive pathogen of the beta herpesvirus family, infecting the majority of the world population^1^. Like all herpesviruses, HCMV persists throughout the lifetime of the host by establishing latent infection from which the virus can later reactivate, causing life-threatening disease in immunocompromised individuals such as transplant recipients and HIV patients^2,3^.

Members of the family *Herpesviridae* are large double stranded DNA viruses, with the HCMV genome encompassing 235 kb, making it one of the largest known viruses to infect humans^4^. During productive infection, transcription from herpesvirus genomes is accomplished by the host RNA polymerase II and regulated by host and viral proteins, leading to a coordinated viral gene expression cascade resulting in the production of infectious progeny. Traditionally, viral genes are divided into three distinct expression groups, differing with respect to their regulation and kinetics. The transcriptional cascade begins with the immediate-early (IE) genes, which require no new cellular or viral protein synthesis for their expression. Next, early (E) genes, including those encoding proteins necessary for DNA replication, are transcribed. These genes are dependent on the presence of viral IE proteins and independent of viral DNA synthesis. Lastly, late (L) gene expression is dependent on viral DNA synthesis and these genes code for the structural proteins necessary for the assembly of new virions^5^. Thus, herpesvirus gene expression during productive infection is assumed to represent a classic sequential regulatory cascade.

Historically, metabolic inhibitors have been used to classify viral genes into IE, E and L temporal expression profiles. Inhibitors of protein synthesis, such as cycloheximide (CHX), lead to the specific accumulation of IE transcripts and these genes were assumed to be expressed at the onset of viral gene expression. Inhibitors of viral DNA replication, such as phosphonoformate (PFA), lead to the specific depletion of transcripts dependent on or enhanced by the onset of viral DNA synthesis and these genes were assumed to be expressed with late kinetics. Early studies provided the framework for mapping viral gene expression kinetics using hybridization methodologies^6–8^ and later on by using microarrays^9^. The reannotation of the HCMV transcriptional landscape has elucidated the crowded nature of its genome and its ability to encode many overlapping RNAs^4,10^ but HCMV gene expression dynamics has not been analysed using unbiased transcriptomic methods.

Viral gene expression during latent infection has been extremely hard to define and is still a matter of controversy^11^. In latently infected cells, the viral genome is maintained in a repressed state where no viral DNA synthesis occurs and there is no production of new infectious virions^12,13^. Since the major immediate-early promoter (MIEP) drives the expression of IE1 and IE2, two potent transactivators of viral gene expression, its regulation has been extensively studied^14-15^ and it is well established that it is chromatinized and repressed in hematopoietic cells, in which latency is established^13,16,17^. Thus, repression of the MIEP is considered a major mechanism dictating latent infection. However, viral gene expression is generally repressed in latent infection, and whether the MIEP is uniquely regulated has not been addressed in a thorough manner. In parallel to studies that focused on understanding the repression of the MIEP, there were extensive efforts to decipher the latent HCMV transcriptome. The leading idea for many years was that in latency, viral gene expression is generally silenced and only a limited number of genes—the presumed latency genes—are expressed^18,19^. However, recent transcriptomic studies on latent hematopoietic cells, including research from our lab, revealed that viral genes are broadly expressed, but at low levels^20,21^. These findings have complicated the characterization of latent viral gene expression and its distinction from lytic viral gene expression as well as the understanding of how the transcriptome reflects latency regulation.

Using transcriptome wide sequencing, we perform a dense time course along both lytic and latent HCMV infections. We define kinetic classes of HCMV transcripts along productive infection, revealing novel patterns of viral gene expression. We uncover that the sensitivity of viral transcripts to translation and replication inhibitors and their temporal dynamics along infection, are often two independent properties, implying that expression of many viral genes is dictated by multiple regulation modules. In latent infection, we show that at early time points post infection the latent transcriptome, in experimental models, is highly dominated by virion-associated input RNA. By unbiasedly measuring the effects of a wide array of chromatin modifiers that enhance viral gene expression, we reveal that the characteristic of latent cells, in terms of viral gene expression, is distinctive repression of IE genes. These findings suggest that the historical terms which defined the basic herpesvirus expression cascade during lytic infection do not capture the full complexity of herpesvirus gene expression dynamics and that latency is characterized by broad repression of viral gene expression with an additional, unique repression of IE genes.

## Results

### Temporal analysis of HCMV gene expression along lytic and latent infections

In order to comprehensively define viral gene expression kinetics along productive infection, and to resolve the molecular events which precede the broad but repressed transcriptional state in latency, we aimed to determine the HCMV temporal gene expression cascade along lytic and latent infections in human foreskin fibroblasts and monocytes, respectively.

Fibroblasts and primary CD14+ monocytes were infected simultaneously with the same TB40 HCMV strain and were harvested for RNA-sequencing (RNA-seq) over the time course of 4, 8, 12, 24, 48, and 72 hours post infection (hpi). In order to determine immediate early genes, cells were treated with CHX, an inhibitor of protein synthesis, and these samples were harvested for RNA-seq at 8 hpi. Additionally, to define true late genes, we treated infected fibroblasts and CD14+ monocytes with PFA, a viral DNA replication inhibitor, and harvested for RNA-seq at 24/48 and 72 hpi, respectively (Figure 1A). Latent infection in CD14+ monocytes was verified by the lack of viral progeny at several times post infection.

**Figure 1.**
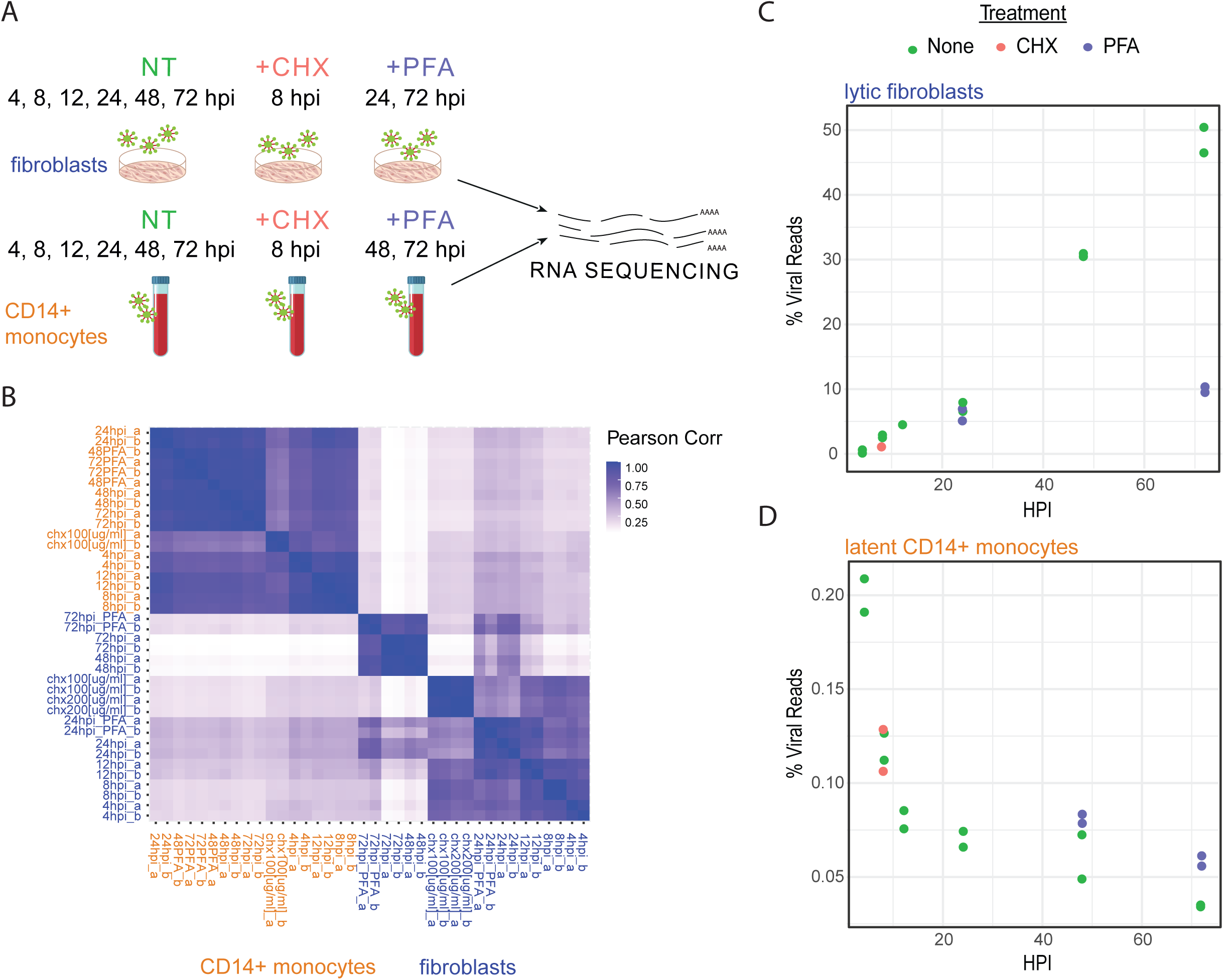
RNA sequencing along HCMV infection in fibroblasts and CD14+ monocytes. (A) Schematic representation of the experimental setup. Fibroblasts and CD14+ monocytes infected with HCMV strain TB40E-GFP were harvested at 4, 8, 12, 24, 48 and 72 hpi for RNA sequencing. In addition, infected fibroblasts and CD14+ monocytes were either treated with Phosphonoformic acid (PFA) and harvested at 24 and 72 or 48 and 72 hpi, respectively or treated with cycloheximide (CHX) and harvested at 8 hpi. All RNA-seq libraries were done in two biological replicates (B) Heatmap depicting the Pearson correlations between all RNA-seq samples based on viral and host reads. Fibroblast samples are marked in blue and CD14+ monocyte samples are marked in orange. All biological replicates are denoted by a/b. (C, D) Percentage of HCMV reads out of total mRNA reads along infection in fibroblasts (C) and CD14+ monocytes (D). The plots include the untreated (green), CHX (red), and PFA (purple) samples with both replicates of each sample.

We prepared two independent biological replicates for all time points, treatments, and cell types. When applying hierarchical clustering on the correlations of host and viral gene expression in each sample, biological replicates grouped together, supporting the reproducibility of our measurements (Figure 1B and Supplementary Fig. S1A). Interestingly, replicates from all time points and treatments in latently infected CD14+ monocytes strongly correlate, indicating there are few changes in the transcriptome along the establishment of latency. In contrast, for infected fibroblasts each time point or treatment represents a more distinct state along productive infection (Figure 1B, Supplementary Fig. S1A and B).

We calculated the percent of HCMV transcripts along infection in the lytic and latent models. In lytic infection, viral gene expression steadily increases between 4 and 72 hours, where viral transcripts compose up to half of the total mRNA by 72 hpi. As expected, in the 72 hpi PFA treated fibroblast samples viral transcripts make up only around 10% of the total mRNA, likely due to less viral DNA template (Figure 1C). In stark contrast, viral transcripts compose less than 0.25% of the total polyadenylated RNA in latently infected monocytes and the percentage of viral transcripts continuously decreases from the earliest points of monocyte infection through 72 hpi. Surprisingly, PFA treated latently infected monocytes expressed slightly higher levels of viral genes likely due to changes in the host cell upon drug treatment (Figure 1D).

### HCMV gene expression kinetics along lytic infection reveals complex, multi-faced regulation

We first analyzed viral gene expression kinetics in lytically infected fibroblasts. After filtering for minimal expression, we were able to reliably quantify the majority of viral genes (134 genes). For each viral gene, we calculated its relative expression pattern out of total viral reads in a given time point or treatment. We performed hierarchical clustering based on the normalized viral gene expression, dividing viral genes into seven distinct temporal classes (Figure 2A). We term these clusters TC1 to TC7 (Figure 2B, Table S1). There was overall good correspondence between classical temporal classes and our measurements. For example, the two canonical immediate early genes UL123 and UL122^22^ were classified as TC1 and were already highly expressed at 4 hpi and their expression was not affected by translation inhibition with CHX. The genes in clusters TC2 and TC3 have early and delayed early kinetics, and contain many genes classically defined as early. The late clusters (TC5-TC7) contain many genes traditionally defined as late. The known late genes UL94^23^, UL75^24,25^, and UL32^26^ were classified as TC7 which showed late kinetics (peak expression 48-72 hpi) and their expression was diminished when viral DNA replication was inhibited by PFA (Figure 2A). In general, there was high agreement between the mRNA temporal classes we defined and previously determined protein temporal classifications ^27^ (Figure 2A lower panel).

**Figure 2.**
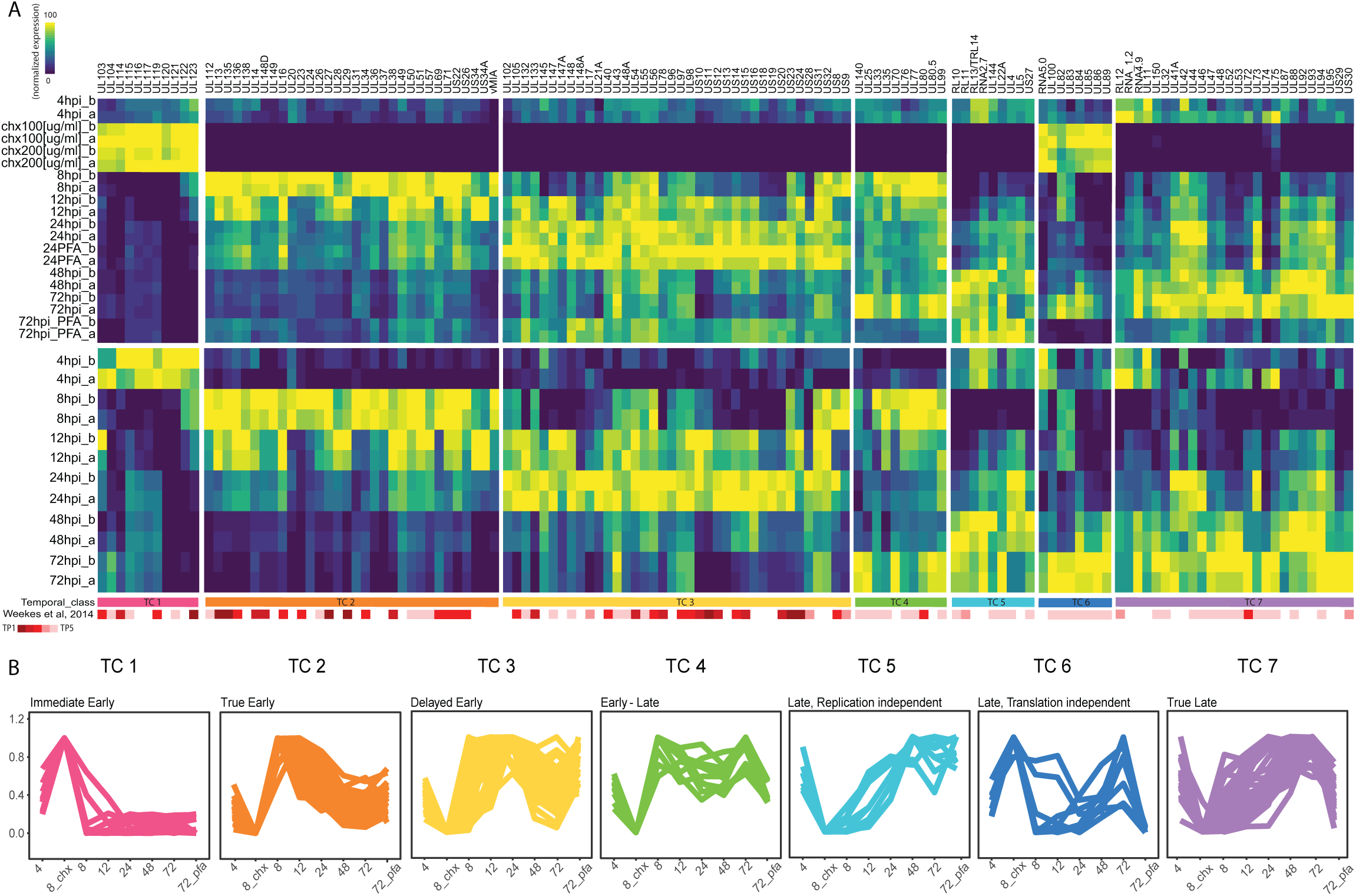
Temporal Classes of HCMV genes along infection of fibroblasts. (A) Heatmap depicting normalized relative expression levels of viral transcripts. Viral genes were clustered into seven temporal classes based on their expression pattern along time, as well as following CHX and PFA treatments. Scaled normalized expression patterns of viral genes are shown including CHX and PFA samples (upper panel) and without the CHX and PFA samples (lower panel). The bars below the panels show the annotations according to our defined temporal classes (top bar) and according to the protein temporal classes that were defined by Weekes et al._10_ (B) Expression profiles of all viral genes for each temporal class (TC) along infection of fibroblasts, averaged across replicates with description for each TC. Values are normalized to the maximal expression of each gene.

It is important to note that the seven TCs reflect how each viral gene is expressed relative to all other viral genes in the same sample. Therefore, the data analysis represents unique expression patterns. Looking at the normalized expression levels of viral genes along infection in RPKM (reads per kilobase per million mapped reads), which more closely reflects their absolute quantities, almost all viral genes are most highly expressed at 72 hpi regardless of the TC in which they were classified (Supplementary Fig. S2). This is expected due to viral DNA replication that leads to high levels of template DNA and a corresponding increase in viral transcription levels at later stages of infection. A unique outlier is UL123 (encoding for IE1) whose highest expression was measured at 8 hpi, and in absolute terms it was hardly expressed at 72 hpi (Supplementary Fig. S2). This unique expression pattern points to the critical role IE1 plays at early stages of the HCMV life cycle and to distinct regulation it is subjected to^28^.

Importantly, we noticed that for numerous viral genes the temporal expression along the course of infection and the expression under CHX and PFA treatments do not fully align, revealing more than one regulatory mode of expression for these genes. For example, TC4 is composed of genes that show early expression kinetics with high expression at 8 hpi. However, these genes also show dependency on DNA replication because PFA treatment greatly diminishes their expression and consequently, they demonstrate a second wave of expression at 72 hpi. Interestingly viral genes that are considered canonical late genes such as UL99 (encoding for pp28) and UL80 (encoding for capsid scaffold protein) are part of this temporal class. TC5 represents a group of genes with late expression kinetics which resembles the kinetics of TC6 and TC7, but the relative expression of genes in TC5 did not depend on DNA replication and was unaffected by PFA treatment (Figure 2A, upper panel). Furthermore, genes in TC6 which includes UL83 (encoding for the major tegument protein pp65)^29,30^ and UL86 (encoding for the major capsid protein)^31^ exhibit late kinetics and dependency on DNA replication, with their highest relative expression at 48-72 hpi and this expression is drastically reduced upon PFA treatment. Surprisingly, however, these genes also exhibit expression that is independent of de novo protein synthesis upon CHX treatment (Figure 2A, upper panel). These results reveal a clear deviation from the classical definition of the herpesvirus gene expression cascade which traditionally relies on the dependence of protein synthesis and sensitivity of viral genes to viral DNA replication in order to define IE and L genes. The canonical naming is oversimplified and even misleading; for numerous viral genes there are several modules that regulate their expression. We therefore propose a division in to seven temporal classes that incorporates the canonical nomenclature but better reflects the complexity of viral gene expression: TC1: Immediate Early, TC2-Early, TC3-Delayed early, TC4: Early-late, TC5-Late-replication independent, TC6-Late-translation independent, TC7-Late (Figure 2B).

### A group of late HCMV transcripts is also expressed in a protein synthesis – independent manner

Analyzing immediate early gene expression in the presence of CHX can be hindered by the possible existence of virion-associated input RNA nonspecifically bound by or packaged within the virus particles and delivered to newly infected cells^32^. Indeed, many of the transcripts that presented late expression kinetics (TC5-TC7) are also relatively highly expressed at 4 hpi. Since TC6 presented such an unexpected expression profile composed of transcripts with late kinetics as well as expression that is independent of protein synthesis, we wanted to rule out the possibility that these transcripts might represent unique, stable RNA species which entered the infected cell with the virus as “input” RNA and remained intact for 8 hours, the time point at which we harvested the CHX samples. In order to assess if the expression of these genes at 8 hpi is independent of transcription, HCMV infected fibroblasts were treated with actinomycin D (ActD), an RNA polymerase inhibitor. Using real time quantitative PCR analysis for TC6 transcripts, we show that the expression of these transcripts, such as UL83 and UL84, is abolished with ActD treatment, indicating our measurements at 8 hpi represent true viral RNA transcription occurring in the newly infected cells (Figure 3A). We also repeated this experiment and analyzed the expression of these genes relative to total viral gene expression at different time points following CHX treatment. Indeed, we detected a higher relative expression of TC6 transcripts (UL83 and UL84) upon CHX treatment, whereas the late transcripts UL32 and UL99 did not show an increase in their expression (Figure 3B). Overall, these results illustrate de novo expression of traditional late genes in a protein synthesis independent manner and provide further support to the validity of the new temporal classes we have identified.

**Figure 3.**
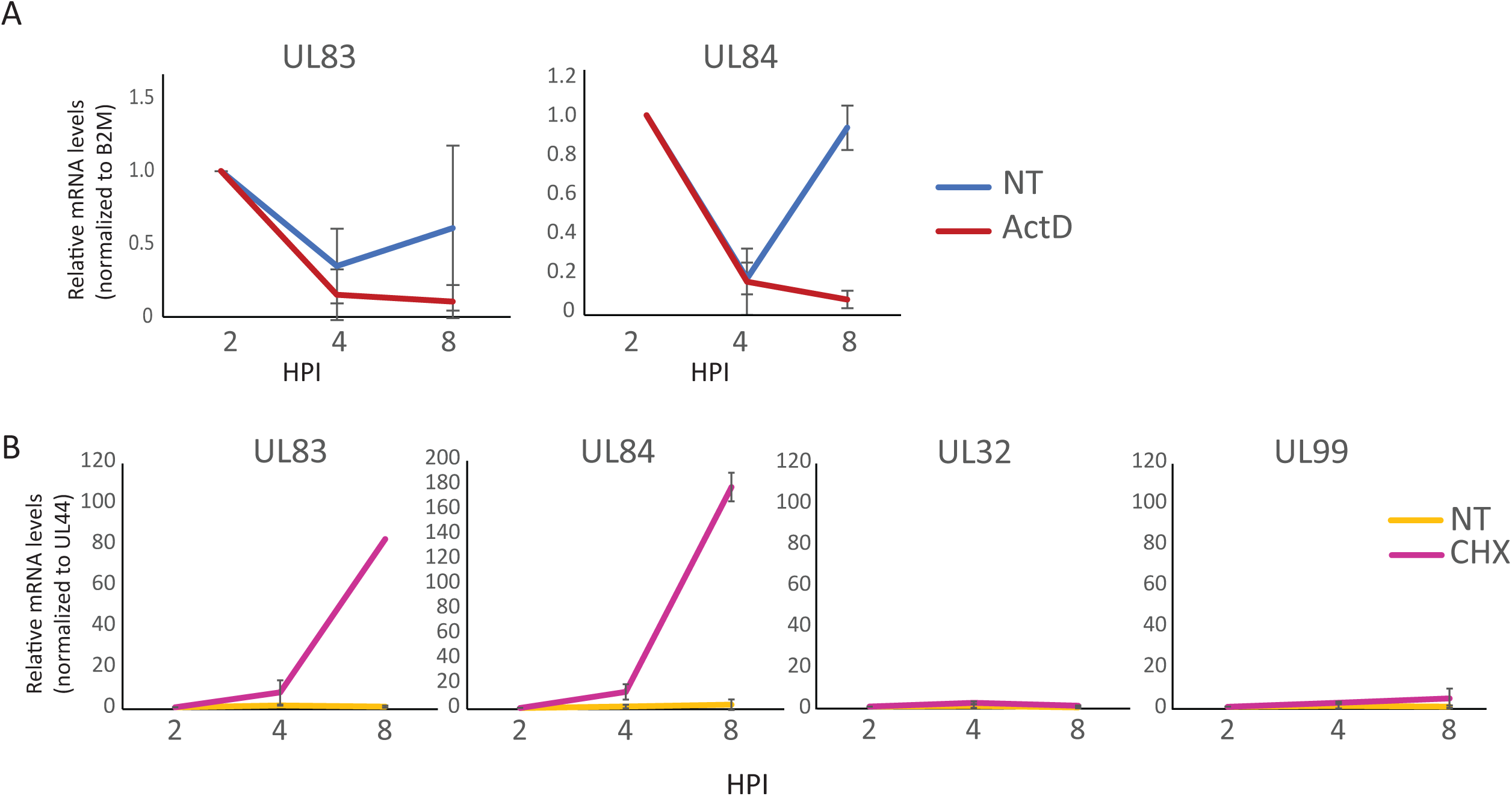
Translation-independent transcription of a group of late transcripts. (A) Fibroblasts were infected with HCMV and were then treated with actinomycinD (actD) or left untreated (NT) at 2 hpi. mRNA levels of the viral transcripts UL83 and UL84 (TC6) were quantified by qRT-PCR at 2, 4, and 8 hpi corresponding to 0, and 2, and 6 hours post actD treatment. RNA levels are normalized to the host B2M transcript. Mean and standard deviation are shown. (B) Fibroblasts were infected with HCMV and cells were treated with CHX or left untreated (NT) at 2 hpi. mRNA levels of viral transcripts UL83, UL84, UL32, and UL99 were quantified by qRT-PCR at 2, 4, and 8 hpi. RNA levels are normalized to viral transcript UL44. Mean and standard deviation are shown.

### High levels of input RNA at 4hpi

High levels of late transcripts in the RNA-seq samples at 4 hpi suggested a portion of the transcriptome at 4 hpi likely originates from virion-derived RNA, rather than RNA that had been synthesized following infection ^32^. In order to better distinguish virion-associated input RNA from genuine transcription, we performed RNA-seq of both HCMV infected human fibroblasts and CD14+ monocytes at 1-hour post infection, a time point which likely precedes the onset of expression for most viral genes. The expression profiles between 1 and 4 hours post infection in both systems highly correlate with each other (Figure 4A), indicating the significant presence of input RNA even at 4 hours post infection. However, in lytic infection (Figure 4A, left panel) TC1 genes, which are expressed with immediate early kinetics, show significantly higher expression by 4 hpi, illustrating clear transcription of this set of genes above the background noise of input RNA. In contrast, in CD14+ monocytes there is no increased expression of specific transcripts at 4 hpi compared to 1 hpi. If transcription of viral IE genes occurs at 4 hpi, it is low and masked by a relatively high level of input RNA (Figure 4A, right panel). We examined whether the lack of increased transcript levels at early time points post infection, together with the gradual decrease in the percent of HCMV transcripts along infection in monocytes (Figure 1D), might indicate large quantities of input RNA and no de-novo viral transcription in this system. Calculation of the decay rate of viral mRNA levels across infection in the infected CD14+ monocytes suggests that by 12 hpi significant portions of the RNA originate from low levels of RNA synthesis (Figure 4B). The source of viral reads in CD14+ monocytes is initially dominated by the decaying input RNA but early in infection there is onset of low-level RNA synthesis, which eventually dominates the measured viral transcripts.

**Figure 4.**
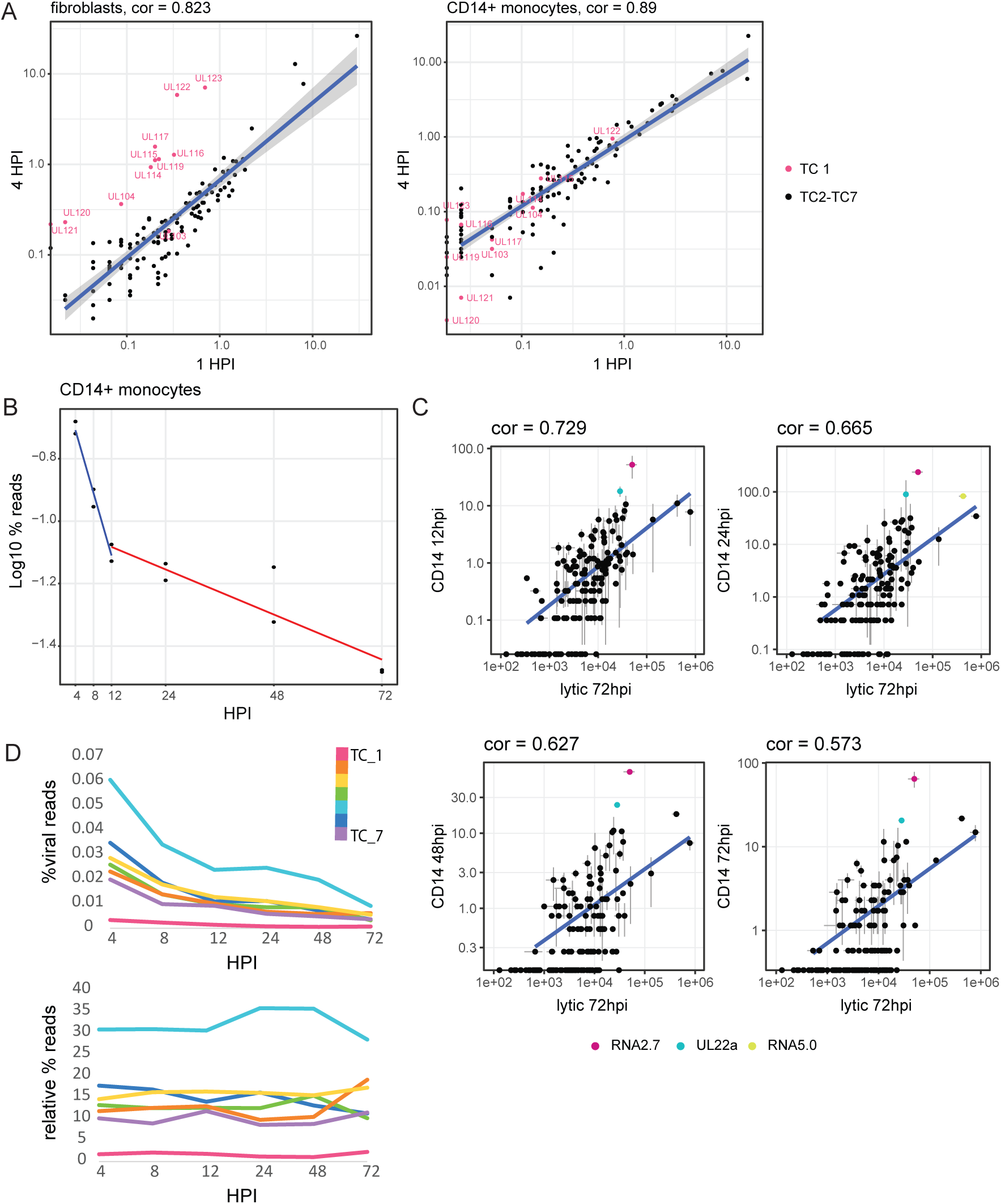
Low level transcription in HCMV infected CD14+ monocytes. (A) Scatter plot showing the read number for each viral gene at 4 hpi versus 1 hpi in fibroblasts (left panel) and CD14+ monocytes (right panel). Viral immediate early genes (TC 1) are marked in pink. Spearman correlations are indicated. (B) Percentage of viral reads out of total mRNA reads along HCMV infection of CD14+ monocytes. The calculated overall decay (half-life) of viral transcripts at 1−12 hpi (0.68 hours) is denoted by a blue regression line and at 12−72 hpi (8.93 hours) is denoted by a red regression line. (C) Scatter plots showing the normalized read number for viral genes at 72 hpi in fibroblasts versus 12, 24, 48, and 72 hpi in CD14+ monocytes. Spearman correlations are indicated. Labeled in color are viral transcripts whose expression significantly deviated (p.value<0.05) from the correlation (D) Expression Profile of all temporal classes (TCs) along HCMV infection of CD14+ monocytes, as calculated by percentage of reads from all viral genes in a TC out of all reads (top) or out of all viral reads (bottom). Mean values of replicates are presented.

### Latently infected CD14+ monocytes express genes that resemble the late lytic profile throughout infection

Although the latency transcriptome profile at early time points post infection was masked by input RNA, we examined if we could capture differential expression of our newly defined TCs along later time points in latent infection. We compared the expression profiles measured at 12, 24, 48 and 72 hpi in CD14+ monocytes to the expression we measured in late lytic infection. We obtained a significant correlation between viral gene expression in infected monocytes with the late lytic expression in fibroblasts and the correlation coefficient dropped over time due to the gradual reduction in the number of viral reads (Figure 4C). The only genes whose expression was significantly higher in monocytes compared to their relative abundance in fibroblasts at all time points were RNA2.7 and UL22a, both of which are highly expressed transcripts, and their relative high levels are likely related to their high mRNA stability. We further analyzed the expression kinetics of the seven temporal classes of gene groups we defined for lytic infection in the CD14+ data set. As expected from the general decline in viral reads in latent monocytes, the expression of all TC classes declined with time (Figure 4D, upper panel). In contrast to the temporal dynamic which is clearly evident in lytic cells (Supplementary Fig. S3), the proportion of reads from each TC remains relatively constant throughout the course of latent infection, (Figure 4D, lower panel) implying little to no changes in the overall expression profile. This fixed profile likely results from residual input RNA in combination with low-level transcription which may reflect inherent susceptibilities of different loci of the viral genome to the host transcription machinery.

### Diverse epigenetic inhibitors induce viral gene expression in infected monocytes

The cellular environment seems to be a key factor in determining the outcome of HCMV infection. Epigenetic regulation plays an important role in latent infection, where the viral genome is chromatinized and maintained as a repressed episome^33,34^. To verify that the expression we captured in latently infected CD14+ monocytes at 72 hpi indeed reflects low-level transcription, we asked if we could enhance viral gene expression in infected CD14+ monocytes by interfering with host chromatin modifying factors. If so, we would expect to see increased expression of specific viral genes, with variable effects between drugs that target different gene silencing pathways.

To probe a large range of molecules, we performed a high-throughput inhibitor screen for epigenetic regulators that affect HCMV gene expression. Primary CD14+ monocytes were latently infected with an HCMV strain TB40E virus encoding a green fluorescent protein (GFP) tag under a simian virus 40 (SV40) promoter (TB40-GFP)^35,36^. 48 hours post infection, at a time when much of the overload of input RNA has degraded and viral gene expression largely reflects low-level latent transcription, we incubated these cells with 140 different small molecule epigenetic inhibitors, in biological replicates. The inhibitors we tested fell within a broad range of categories, including inhibitors of bromodomains, histone deacetylases, histone methyltransferases, histone demethylases, and others (Figure 5A). The final concentration we used in this screen was set to 1uM based on calibration of six different drugs targeting different functional categories with the aim to reduce drug cytotoxicity (Supplementary Fig. S4). 24 hours following drug treatment (a time point we presumed is likely to reflect the immediate changes due to the drugs on viral gene expression), we screened by flow cytometry for bulk changes in GFP expression as a proxy for changes in viral gene expression compared to a DMSO treated control. For the vast majority of epigenetic inhibitors, biological replicates showed little variation (Figure 5B) and treatments induced little to no changes in the GFP expression. The 15 drugs that fell within the top ten percent for fold change in GFP expression (Figure 5C and 5D) were enriched in HDACs such as Trichostatin A (TSA)^37-38^, Sodium Butyrate^39,40^ and MC1568^41^, and also included the Sirtuin inhibitor AGK2^42-43^ all of which have been reported to enhance herpesvirus gene expression. Additionally, these drugs included UNC0631, histone methyltransferase inhibitor, and Etoposide and Mirin, inhibitors of the DNA damage pathway^44^ (Figure 5E).

**Figure 5.**
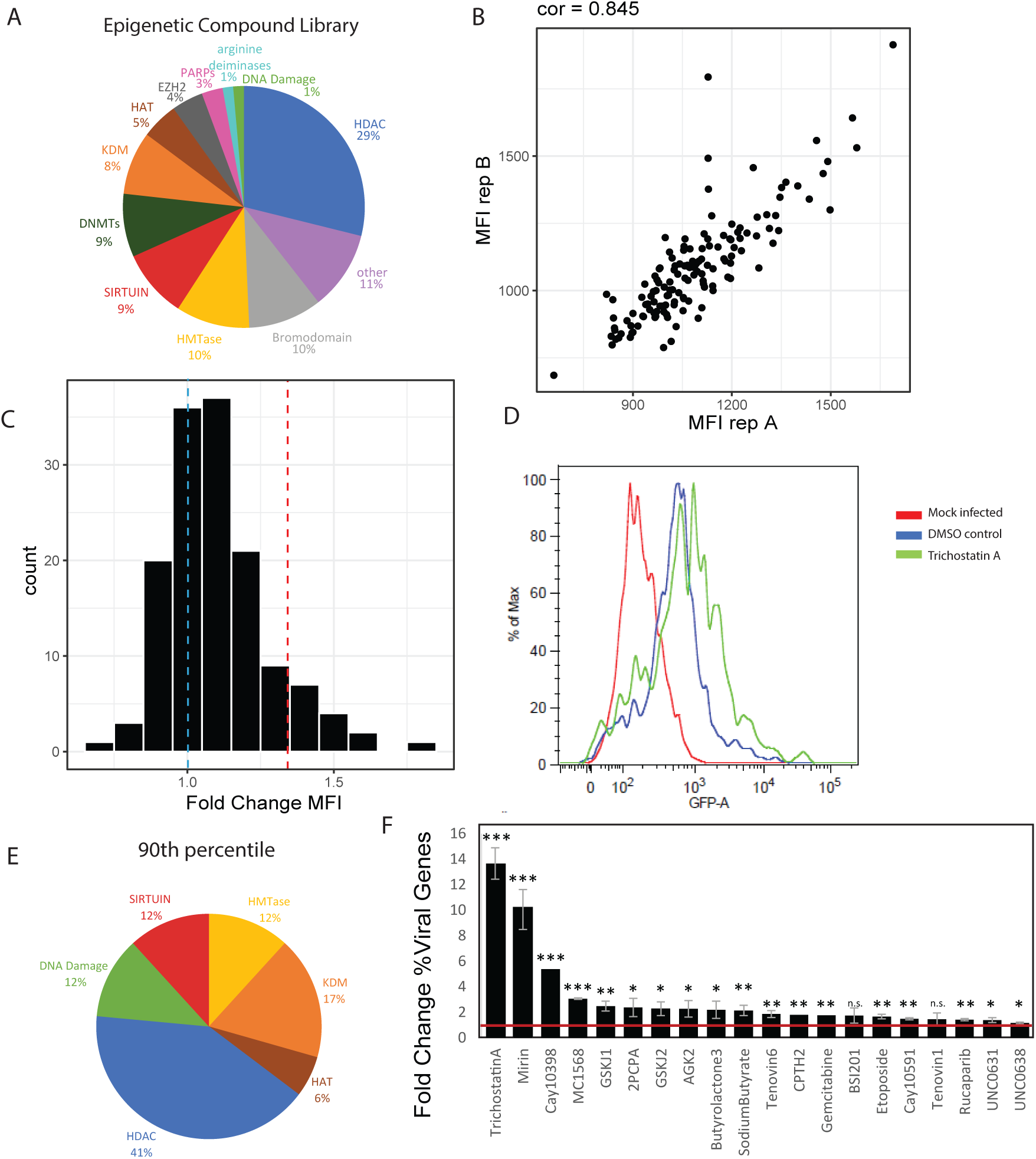
Epigenetic inhibitors induce viral gene transcription in HCMV infected CD14+ monocytes. (A-F) CD14+ monocytes infected with HCMV expressing a GFP reporter were incubated with 1 μM of each of 150 compounds from an epigenetic inhibitor library, for 24 hours at 48 hpi, in biological replicates. (A) A pie chart showing the distribution of drug targets in the epigenetic inhibitor library. (B) Scatter plot showing GFP Mean fluorescence intensity (MFI) values measured by flow cytometry for each treatment of biological replicates. Spearman correlation is indicated. (C) Histogram depicting the distribution of MFI fold change, compared to a DMSO control, for all tested inhibitors. The blue dashed line shows the DMSO control sample (FC=1) and the red dashed line pinpoints the 90th percentile. (D) Flow cytometry analysis of GFP levels for a representative inhibitor, TrichostatinA, whose MFI fold change lies within the 90th percentile, compared to the DMSO sample and uninfected cells. (E) A pie chart showing the distribution of drug targets of the inhibitors in the 90th percentile of MFI fold change. (F) RNA-sequencing was performed in duplicates on HCMV infected CD14+ monocytes treated with 20 different inhibitors and DMSO control. Plot showing fold change of percent viral reads for each treatment compared to DMSO control. Mean and standard deviation of duplicates are presented. Red line marks fold change of 1. p-values representing significant increase in viral gene expression were calculated with t-test ***pval≤0.001, **pval≤0.01, *pval≤0.05, n.s. pval>0.05.

Since the GFP we used in our screen was driven by an exogenous promoter, we next verified that the epigenetic drugs identified in our screen based on GFP expression indeed correspond to the elevation in the expression of additional viral genes. To this end, we performed transcriptome analysis of infected CD14+ monocytes following treatment with 20 drugs from the screen, most with biological replicates. The drugs that were used included the top 10% hits of our flow cytometry based screen as well as several drugs from the top 20% hits that fall into additional functional categories such as poly-ADP ribose polymerase inhibitors. Reassuringly, treatment with most of the inhibitors elicited significant increase in total viral gene expression compared to the DMSO treated control (Figure 5F), with Trichostatin A and Cay10398, two HDAC inhibitors and Mirin, an ATM pathway inhibitor inducing the most significant increases in viral gene expression (13.6-fold, 5.5-fold and 10.3-fold correspondingly).

### Transcription induction in infected monocytes reveals that the hallmark of HCMV latency is distinctive repression of IE genes

To examine which viral genes were affected by each of these drugs, we compared the gene expression profiles of HCMV infected monocytes treated with epigenetic drugs to the control sample. Importantly, viral gene expression following all drug treatments, regardless of their specific target and the levels of viral gene induction, was highly correlated with the control sample (Figure 6A). Statistical analysis showed almost no viral genes that were distinctively induced by each of these drugs. This overall resemblance in viral gene expression between untreated monocytes and monocytes treated with diverse drugs that work by different mechanisms points to two critical conclusions: 1. Although there are substantial amounts of input RNA upon infection, viral RNA expression at 72 hpi mostly reflects low level transcription that can be further enhanced. 2. Upon latent infection of monocytes viral gene expression is repressed and this repression is largely consistent throughout the viral genome, such that treatment with diverse chromatin modifiers results in a uniformly enhanced expression profile.

**Figure 6.**
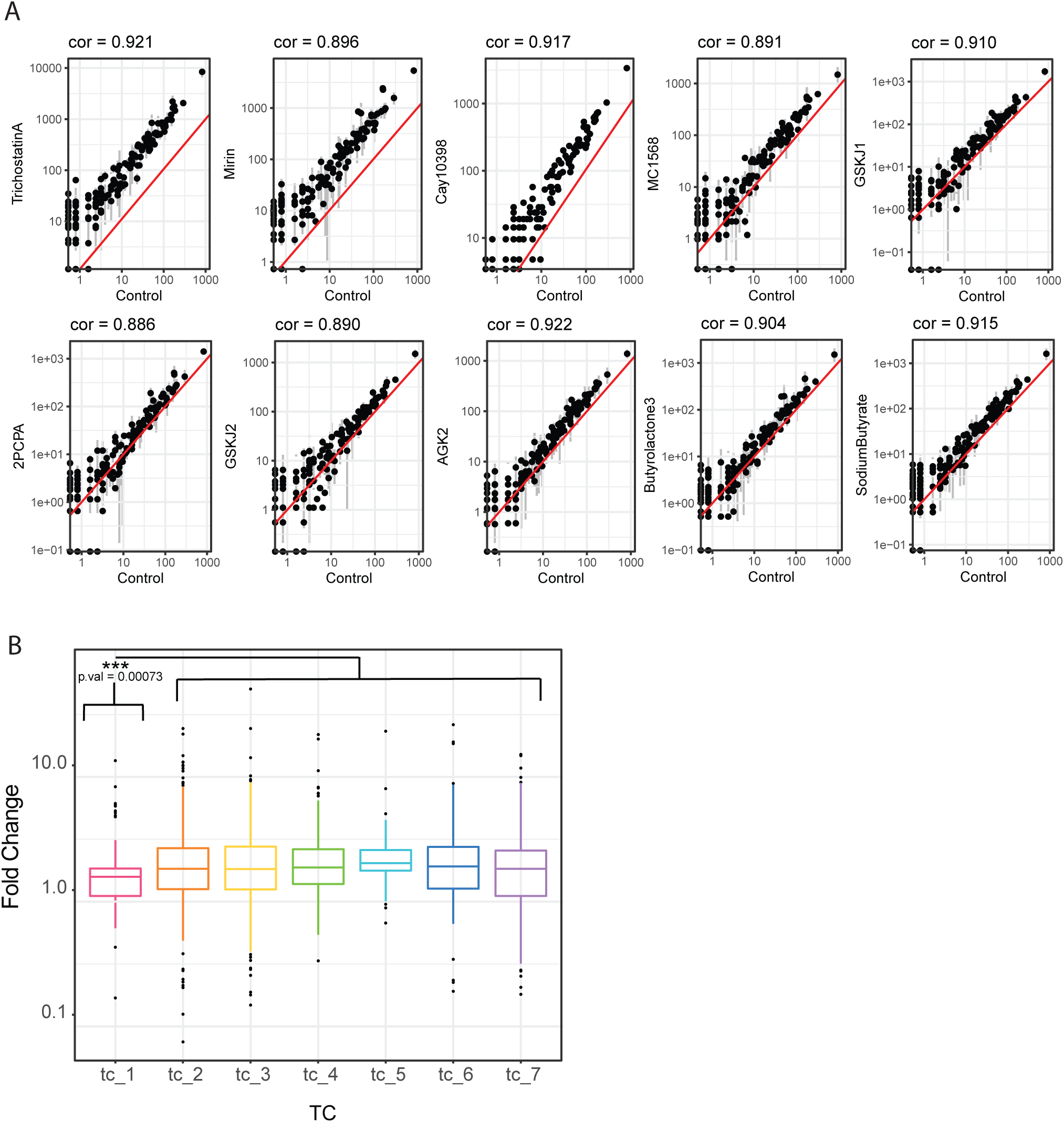
Unique repression of IE genes in HCMV infected CD14+ monocytes. (A) Scatter plots showing the read number for viral genes in HCMV infected CD14+ monocytes treated with DMSO as control versus treatment with ten different inhibitors that elicited the strongest increase in viral gene expression. Spearman correlations are indicated. The red line marks 1:1 ratio. Only in the mirin sample, 2 viral genes (RNA5.0 and RL1) significantly deviated (p.value<=0.05 and FC=>2) from the correlation with the DMSO control (B) Boxplot showing the transcript level fold change of viral class between inhibitors treated and control treated HCMV infected CD14+ monocytes. p-value calculated by t-test comparing TC1 and all other TCs is indicated.

Since viral gene expression is overall very low (between 0.4 to 4% of expressed mRNAs are viral), small but significant changes in a limited number of genes might be difficult to detect and quantify. Therefore, we also analyzed the changes in expression of the temporal classes across the different drugs. This pooled analysis revealed an increase of all TCs (Figure 6B), but interestingly, a common signature of most drugs was the relatively low induction of TC1 expression (IE genes) (Supplementary Fig. S5), and this effect was statistically significant when analyzed across all drugs (Figure 6B). This finding indicates that a defining property of the latent transcriptome is the unique repression of IE transcripts that differs from the general repression of all other viral genes.

To verify that the low induction of TC1 genes is not related to the time point post drug treatment that we analyzed, we performed gene expression analysis at different time points after drug treatment. In this experiment, we focused on TSA, which led to the most significant increase in viral gene expression, thereby providing us with a maximal dynamic range. 48 hpi, CD14+ monocytes were treated with TSA or DMSO as a control. Cells were harvested in replicates for RNA-seq at 0, 4, 12 and 24h post TSA treatment. We observed a small (1.2-fold) but significant increase in viral expression already at 4h post TSA treatment and the induction in viral gene expression increased with time, reaching 14-fold by 24h post treatment (Figure 7A). In agreement with our previous observations, viral gene expression patterns in infected monocytes treated with TSA were highly correlated with the untreated samples at all time points post treatment (Figure 7B). These measurements illustrate that also in the early time points post treatment, TSA does not lead to dynamic changes in viral gene expression but rather to enhancement of the same viral transcription profile that exists in infected monocytes. We next grouped viral transcripts according to their TC and analyzed the induction of different TC gene groups across the time points. Across all time points, viral genes from all TCs increase. However, at 24 hours post TSA treatment, when viral gene induction was the strongest, expression of TC1 genes was induced to a significantly less extent than genes from other TCs (Pval<0.001, Figure 7C). This analysis shows that the induction of viral gene expression in infected monocytes by epigenetic modifiers is associated with the relative unresponsiveness of IE genes, indicating an inherent difference in the repression or activation mechanism between IE genes and other genes during latent infection. As has been previously demonstrated^45^, we found that Trichostatin A, which also caused the highest induction of viral RNA expression, was the only drug that reproducibly led to reactivation from latency in a low proportion of infected monocytes. Overall, these results demonstrate that diverse epigenetic drugs lead to a broad increase in viral gene expression with weaker induction of IE genes which appear to be under additional repressive regulation in latent cells. Nevertheless, when viral genes and IE genes are induced beyond a certain threshold the virus can still reactivate.

**Figure 7.**
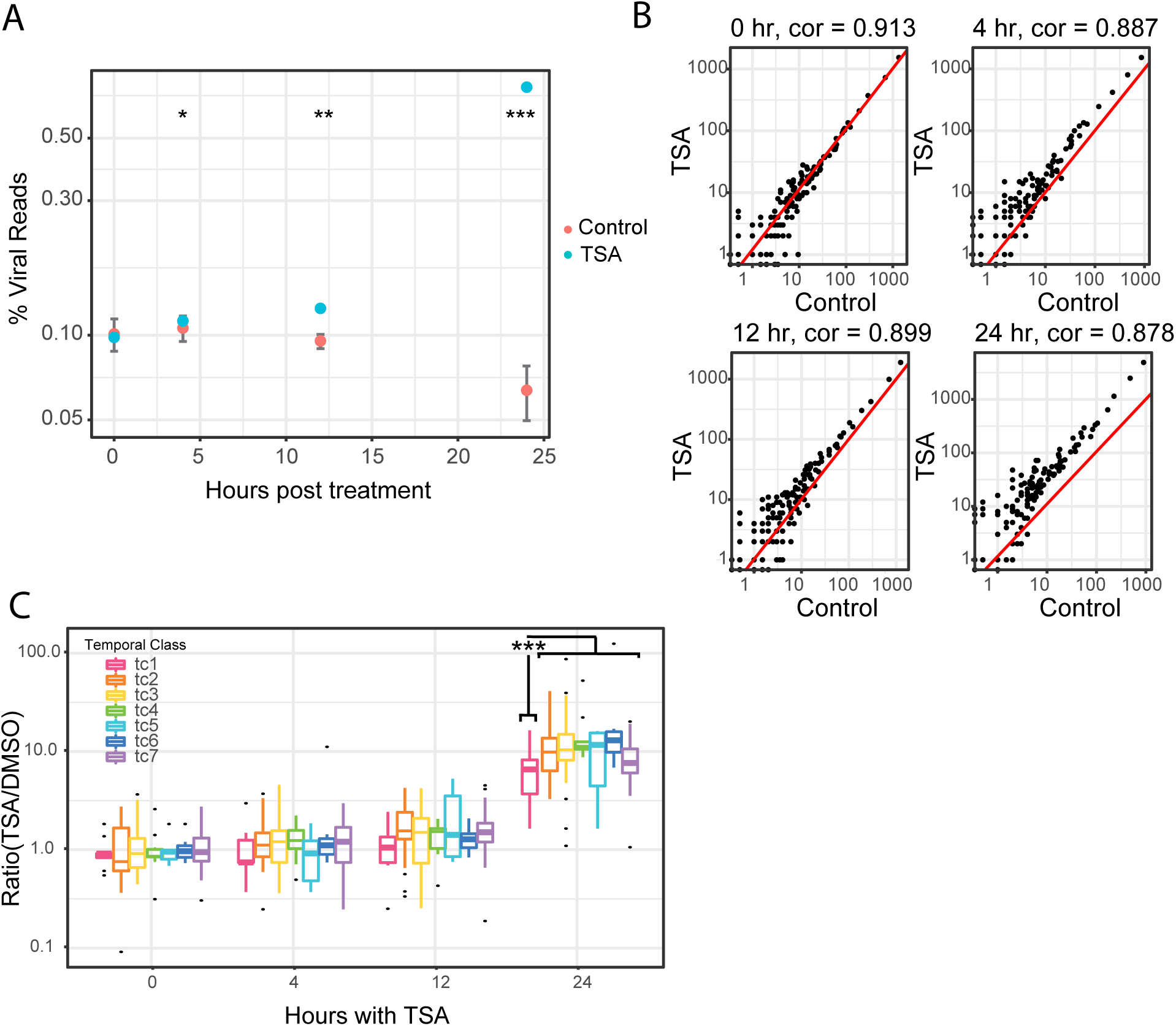
Unique repression of IE genes occurs at different times following TSA treatment. (A-C) RNA-sequencing was performed on HCMV infected CD14+ monocytes treated with TSA and DMSO control, in biological replicates, at 0, 4, 12 and 24 hours after treatment. (A) Percentage of HCMV reads out of total mRNA reads at different time points following treatment. Error bars represent standard deviation in the control samples. p-values were calculated with t test. *pval≤0.05, **pval≤0.01, ***pval≤0.001. (B) Scatter plots showing the read number of viral genes in HCMV infected CD14+ monocytes treated with DMSO control versus TSA treatment at 0, 4, 12 and 24 hours after treatment. Spearman correlations are indicated. The red line marks 1:1 ratio. (C) Boxplot showing the transcript level fold change of viral genes from each temporal class (TC) between TSA and DMSO control treated HCMV infected CD14+ monocytes at different time points following treatment. p-value = 0.0011 calculated by t-test comparing TC1 and all other TCs at 24 hours following treatment is indicated.

### HCMV gene expression induction in latent monocytes leads to reduction in the expression of immune related genes

To further probe the effects of these diverse epigenetic drugs, we analyzed the host gene expression landscapes following treatment with epigenetic modifiers. Gene set enrichment analysis (GSEA) shows that compared to untreated cells, the drug-treated infected monocytes (in which viral gene expression was induced by more than 2-fold) display significant reduction in many immune response related pathways. These include TNFa signaling, IFN response and inflammatory response as well as reduction in pathways related to cell cycle progression and apoptosis (Figure 8A). Remarkably, although the drugs target diverse chromatin modifiers, the changes in host gene expression are common between the different drug treated samples. Monocytes that were treated with BSI201, which did not lead to a significant increase in viral gene expression, showed only marginal reduction in a limited number of these pathways (Figure 8A). Common changes in cellular gene expression likely reflect secondary effects that relate to the increase in viral RNA or protein expression. Furthermore, we observed an inverse correlation between the increase in viral gene expression and the reduction in host gene immune pathways such as IFNa/g responses and TNFa signaling (Figure 8B), supporting the notion that these changes are driven by the increase in viral gene expression. Finally, we examined the kinetics of changes in host gene expression,by analyzing the effects of TSA across the different time points (Supplementary Figure S6). This analysis revealed that down regulation of IFNa and IFNg pathways coincides with the increase in viral gene expression (Figure 8C), further indicating that the enhancement in viral gene expression accounts for a major portion of the changes in cellular gene expression.

**Figure 8.**
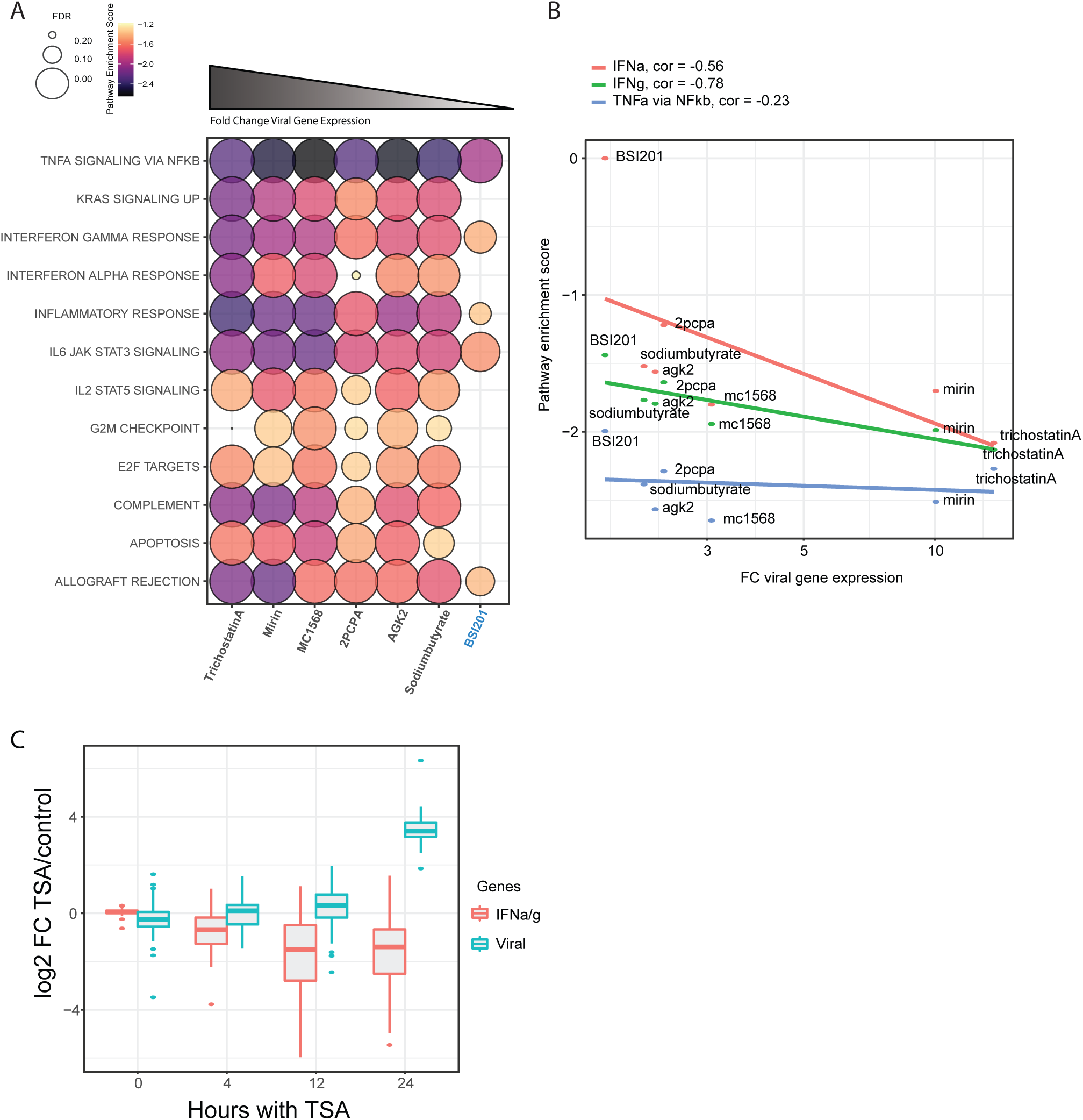
Induction of viral gene expression during HCMV latency leads to reduction in expression of immune related genes. (A). Bubble plot showing enriched human hallmark pathways which are downregulated following inhibitor treatment of infected CD14+ monocytes for drugs which significantly induced more than 1.8-Fold Change in viral gene expression. BSI201, highlighted in blue, did not produce a significant increase in viral gene expression. (B). Scatter plot depicting enrichment scores for three host pathways: interferon alpha response (IFNa, red), interferon gamma response (IFNg, green) and TNF alpha signaling via NFKb (blue) versus viral transcript level fold change for different inhibitor treatments compared to the control. Spearman correlations are indicated. (C) Box plot showing the transcript level log2 Fold change of viral genes and IFNa/g induced host genes in TSA treated samples at 0, 4, 12, and 24 hours after treatment.

## Discussion

Using transcriptome wide sequencing, we performed a dense time course along both lytic and latent HCMV infections. We defined kinetic classes for the majority of HCMV transcripts along productive infection, revealing novel patterns of viral gene expression. We discovered that in contrast to the traditional view of herpesvirus gene expression, the dependency of viral transcripts on metabolic drugs and viral transcripts’ temporal dynamics are often decoupled, implying that expression kinetics and the dependency on protein synthesis or viral DNA replication are frequently two independent properties in viral gene expression regulation. A recent study on alpha herpesvirus, varicella-zoster virus (VZV), which mapped VZV transcripts also examined their expression kinetics and observed unexpected patterns of viral gene expression^46^. In addition to the traditional IE, E and L kinetic classes, they defined groups of genes as Early-Late and Transactivated/True-Late (TA/TL)^46^. In their analysis, Early-Late genes show two waves of expression, one of which is dependent on de novo protein synthesis and a second, later wave of expression which depends on the onset of DNA-replication. These genes are comparable to the HCMV genes characterized here in TC4, which we also name Early-Late. TA/TL genes correspond in unique expression dynamics to our TC6: Late-translation independent gene cluster. These genes show low level expression in the absence of de novo protein production, but their predominant expression occurs at late time points post infection and is highly dependent on viral DNA replication. The authors hypothesized that the production of these transcripts in CHX-treated samples might be due to low-level transactivation by viral tegument proteins delivered with incoming virions. Whether our TC6 genes can be expressed independently of viral proteins or whether they in fact depend on incoming viral tegument proteins is an important question which warrants further research. Regardless, the ability of these genes to be expressed when infection starts suggests they play an additional important role in the early stages of infection or in reactivation.

More globally, in the herpesvirus literature it has become commonplace to use the terms “leaky late” and “true late” to distinguish between genes whose expression is augmented by versus totally dependent on DNA replication, respectively. This terminology has hinted towards a more intricate expression regulation than initially presumed for these viruses. The resemblance between the expression patterns we uncover here to the ones described for VZV,^46^ strongly points to complex expression regulation across diverse herpesviruses. We therefore propose that the clusters we describe, which are based on both regulation and temporal expression, may better capture the multiple modules herpesviruses utilize to tightly regulate their gene expression. One clear implication of these more complex expression patterns is that they do not represent a simple sequential cascade as the traditional IE, E, L model. Therefore, a deeper understanding of herpesvirus gene regulation will be needed to decipher the connectivity between these different expression modules and how they functionally come into play throughout productive infection.

During HCMV latency there is massive repression of viral gene expression. We revealed that at early time points post infection, the latent transcriptome in experimental models is highly dominated by virion-associated input RNA. Looking at viral mRNA decay rates, we can infer that the percent of viral reads in infected CD14+ monocytes is set by the decay of virion associated input RNA together with the onset of low-level RNA synthesis. In order to clearly distinguish newly synthesized mRNAs from background input RNA noise and to unambiguously determine transcription patterns at early time points during latency establishment, it would be necessary to perform metabolic labeling which allows the detection and quantification of newly synthesized RNA species^47–49^. However, since the levels of viral gene expression are incredibly low and metabolic labeling approaches facilitate labeling of only a small portion of the newly synthesized RNA pool, conducting these experiments will be challenging.

When we analyze the relative proportion of reads from each TC in monocytes from 12 hpi and onwards, it remains largely constant. We hypothesize that the latency transcriptome represents overall uniform viral gene repression with default transcription tendencies of the viral genome dictating the low-level transcription profile. To convincingly show this low-level viral gene expression represents true transcription, we screened a wide array of chromatin modifiers and tested their effects on viral gene expression. This screen revealed compounds from several categories that enhanced viral gene expression, including inhibitors of HDACs, Sirtuins and DNA damage response, which were previously implicated in regulation of viral gene expression during herpesvirus latency^44,50–53^. Although these drugs work by diverse mechanisms, causing different changes to the chromatin, characterization of the viral gene expression enhanced by these drugs revealed that they all lead to induction of the same transcriptional program. Importantly, the only distinctive group was TC1-IE genes; induction of IE genes was significantly less prominent than the induction of viral genes from all other TC gene clusters. This implies that repression along the HCMV genome in latent cells is uniform with an additional and unique repression of IE genes. This unique regulation may still rely on epigenetics, but likely involves additional mechanisms. The regulation of IE1 expression by PML-NB proteins in non-permissive cells has been studied by multiple groups with contradicting results ^54,55,56^. We speculate that the absence of a particular transcription factor or as recently suggested, the differing occupancy of polymerase II at the MIEP may be involved ^57,58^. The unique down-regulation of IE genes is well established in the context of lytic infection, where it was shown that repression is mediated through binding of a cis repression sequence and was thought to be important during viral replication.^28^ Our results suggest that in the context of latent infection, a similar or complementary regulatory mechanism might also exist in monocytes. The results from our epigenetic drug screen can also be interpreted to support the “stochastic transcription hypothesis,” which was shown to play a role in MCMV latency and proposes that viral genes become transiently de-silenced in latent viral genomes in a stochastic fashion and not following the canonical temporal cascade of reactivation.^59,60^ Enhanced, broad viral transcription by epigenetic drug treatments can eventually lead to reactivation of latent cells and this may occur once IE gene levels surpass a given threshold.

When analyzing host gene expression due to treatment with our diverse set of drugs, we observed recurrent down regulation of the host innate immune response pathway. In agreement with our previous findings^61^, this indicates that viral gene expression during latent infection does have functional consequences and suggests a constant arms race between viral mRNA or viral protein expression and expression of host innate immune response genes during infection and latency^62^. In summary, our findings suggest that herpesvirus temporal gene expression cascade is dictated by a more intricate set of dependencies than was previously appreciated and that as envisioned for many years, the trademark of gene expression during HCMV latency is the unique repression of IE genes.

## Acknowledgments

We thank Stern-Ginossar lab members, Oren Kobiler and Igor Ulitsky for providing valuable feedback. We thank the Weizmann flow cytometry units for technical assistance. This study was supported by a European Research Council consolidator grant to N.S-G (CoG-2019-864012). N.S-G is an incumbent of the Skirball Career Development Chair in New Scientists and is a member of the European Molecular Biology Organization (EMBO) Young Investigator Program. The authors declare no competing interests.

## Materials and Methods

### Cells and viruses

Human foreskin fibroblasts (ATCC CRL-1634) were grown in Dulbecco’s Modified Eagle’s Medium (DMEM) with 10% heat-inactivated fetal bovine serum (FBS), 2 mM L-glutamine, and 100 units/ml penicillin and streptomycin (Beit-Haemek, Israel) and maintained at 37°C in a 5% CO2 incubator. Primary CD14+ monocytes were isolated from fresh venous blood, obtained from healthy donors, using a Lymphoprep (StemCell Technologies) density gradient followed by magnetically activated cell sorting with CD14 magnetic beads (Miltenyi Biotec). CD14+ cells were cultured in X-Vivo 15 medium (Lonza) supplemented with 2.25 mM L-glutamine and maintained at 37°C in a 5% CO2 incubator. The bacterial artificial chromosome (BAC) containing the clinical strain TB40E^63^ with an SV40-GFP tag (TB40E-GFP) was described previously^35,36^. This strain lacks the US2-US6 region, therefore, these genes were not included in our analysis. Virus was propagated by adenofection of infectious BAC DNA into fibroblasts^64^. Viral stocks were concentrated by ultracentrifugation at 30,000xg at 4°C for 120 min. Infectious virus yields were assayed on fibroblasts and THP1 cells.

### Infection procedures

For the time course analysis fibroblasts and CD14+ monocytes were infected with the same stock of HCMV strain TB40E-GFP at a multiplicity of infection (MOI) of 1 and 10, respectively. For all other experiments CD14+ monocytes were infected with HCMV strain TB40E-GFP at an MOI of 5. Cells were incubated with the virus for 2h, washed, and supplemented with fresh medium. Infection was monitored 2-3 dpi by measurement of GFP expression on a BD accuri flow cytometer and analysis by FlowJo. Substantial shift in GFP levels, seen in most infected fibroblasts is indicative of productive infection, while a small shift in GFP levels, as seen in all CD14+ monocytes indicates that the cells were infected but the virus is repressed, as expected in latent infection.

### Cell treatments

CHX was added to infected fibroblasts and CD14+ monocytes immediately after infection at a final concentration of 100 ug/ml and 200 ug/ml or at 100 ug/ml, respectively. PFA was added to fibroblasts and CD14+ monocytes immediately after infection at a final concentration of 400 ug/ml. ActD was added to infected fibroblasts at a final concentration of 5uM.

### RNA library construction

For RNA-seq time course experiments in fibroblasts and CD14+ monocytes, cells were washed with PBS and then collected with Tri-Reagent (Sigma-Aldrich), total RNA was extracted by phase separation and poly(A) selection was performed using Dynabeads mRNA DIRECT Purification Kit (Invitrogen) according to the manufacturer’s protocol. RNA-seq libraries were generated as previously described^65^. Briefly, mRNA samples of ∼4ng were subjected to DNaseI treatment and 3′ dephosphorylation using FastAP Thermosensitive Alkaline Phosphatase (Thermo Scientific) and T4 PNK (NEB) followed by 3′ adaptor ligation using T4 ligase (NEB). The ligated products were used for reverse transcription with SSIII (Invitrogen) for first-strand cDNA synthesis. The cDNA products were 3′ ligated with a second adaptor using T4 ligase and amplified with 8 cycles of PCR for final library products of 200–300 base pairs. For epigenetic drug-treated CD14+ monocyte samples, RNA libraries were generated from samples of ∼100,000 cells according to the MARS-seq protocol^66^.

### Next-generation sequencing and data analysis

All RNA-Seq libraries (pooled at equimolar concentration) were sequenced using Novaseq6000 (Illumina), with read parameters: Read1: 72 cycles and Read2: 15 cycles. For the time course libraries, raw sequences were first trimmed at their 3′ end, removing the illumina adapter and polyA tail. Alignment was performed using Bowtie (allowing up to 2 mismatches) and reads were aligned to concatenation of the human (hg19) and the viral genomes (NCBI EF999921.1). Reads aligned to ribosomal RNA were removed. Reads that were not aligned to the genome were then aligned to the transcriptome.

Analysis of libraries generated with the MARS-seq protocol was done as previously described^20^. Briefly, 37-bp reads were aligned using Bowtie (allowing up to 2 mismatches) to concatenation of the human (hg19) and the viral genomes (NCBI EF999921.1). Counting of reads per gene was done based on unique molecular identifiers (UMIs) (8 bp). The transcription units of the virus were based on NCBI annotations, with some changes, including merging several transcripts (considering that the library maps only the 3’ ends of transcripts) and adding some antisense transcripts. All analyses and figures were done using in-house R-scripts. Epigenetic drug treated CD14+ monocyte samples which had less than 100,000 UMIs were left out from further analysis leaving three drug treatments without duplicates-Cay10398, CPTH2, and Gemcitabine.

### Inhibitor screen

The compounds used in the epigenetic inhibitor screen were purchased from Cayman Chemical as a complete Epigenetics Screening Library (96-Well, Item No. 11076). To calibrate cytotoxicity, HCMV infected CD14+ monocytes were treated with six inhibitors at increasing concentrations (500nM,1uM,10uM) for 48 hours. Cells were washed, resuspended in PBS and stained with 0.5ug/ml of PI for 10 minutes prior to analysis by FACs-LSRII. The 1*μ*M concentration was chosen for the complete screen as it caused minimal cytotoxicity. For the inhibitor screen at 48 hpi, TB40-infected CD14+ monocytes were divided into 1.2 ml tubes containing ∼100,000 cells per tube and inhibitors or DMSO for negative control were added to a final concentration of 1uM. After 24 hours of incubation, Flow cytometry was performed using an LSRII, and analysis was performed using FlowJo software.

### Differential expression and enrichment analysis

Differential expression analysis was done with DESeq2 (version 1.22.2)^67^ using default parameters, with the number of reads in each of the samples as an input. The normalized number of reads according to DESeq2 were used for enrichment analysis using GSEA (version 4.0.3)^68^. The MSigDB hallmark (version 7.1) gene sets were used^69^. The GSEA plots were created based on the GSEA output with in house R scripts using R package enrichplot.

### Quantitative real-time PCR analysis

For analysis of RNA expression, total RNA was extracted using Tri-Reagent (Sigma) according to the manufacturer’s protocol. cDNA was prepared using the qScript cDNA Synthesis Kit (Quanta Biosciences) according to the manufacturer’s protocol. Real time PCR was performed using the SYBR Green PCR master-mix (ABI) on the QuantStudio 12K Flex (ABI) with the following primers (forward, reverse):

UL123 - (TCCCGCTTATCCTCAGGTACA, TGAGCCTTTCGAGGACATGAA)

UL84 - (TCAAAGCATACGCTGAATCG, GCTTACAGTCTTGCGGTTCC)

UL83 - (CCAGCGTGACGTGCATAAAG, TCGTGTTTCCCACCAAGGAC)

UL44 - (AGCAAGGACCTGACCAAGTT, GCCGAGCTGAACTCCATATT)

UL99 - (GGGAGGATGACGATAACGAG, TGCCGCTACTACTGTCGTTT)

UL32 - (GGTTTCTGGCTCGTGGATGTCG, CACACAACACCGTCGTCCGATTAC)

RNA 2.7 - (TCCTACCTACCACGAATCGC, GTTGGGAATCGTCGACTTTG)

B2M - (TGCTGTCTCCATGTTTGATGTATCT, TCTCTGCTCCCCACCTCTAAGT)

---

**Fig S1.**
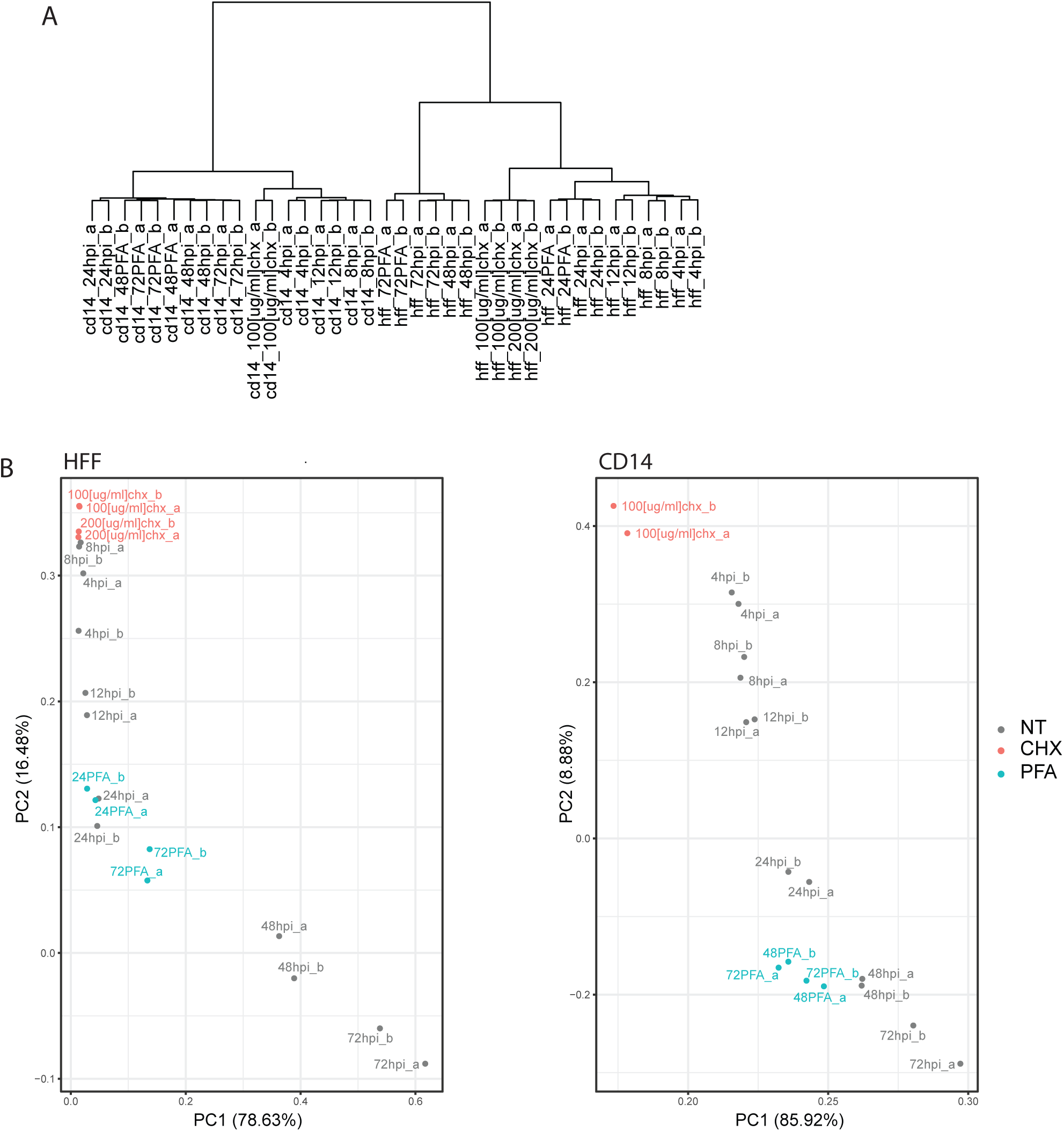
RNA sequencing along HCMV infection in fibroblasts and CD14+ monocytes. (A) Hierarchical clustering dendrogram of all samples. Cell lines cluster separately, and biological replicates cluster together. (B) PCA analysis for each data set (HFF and CD14+) based on host and viral transcript reads. CHX treated samples are highlighted in red and PFA treated samples in blue.

**Fig S2.**
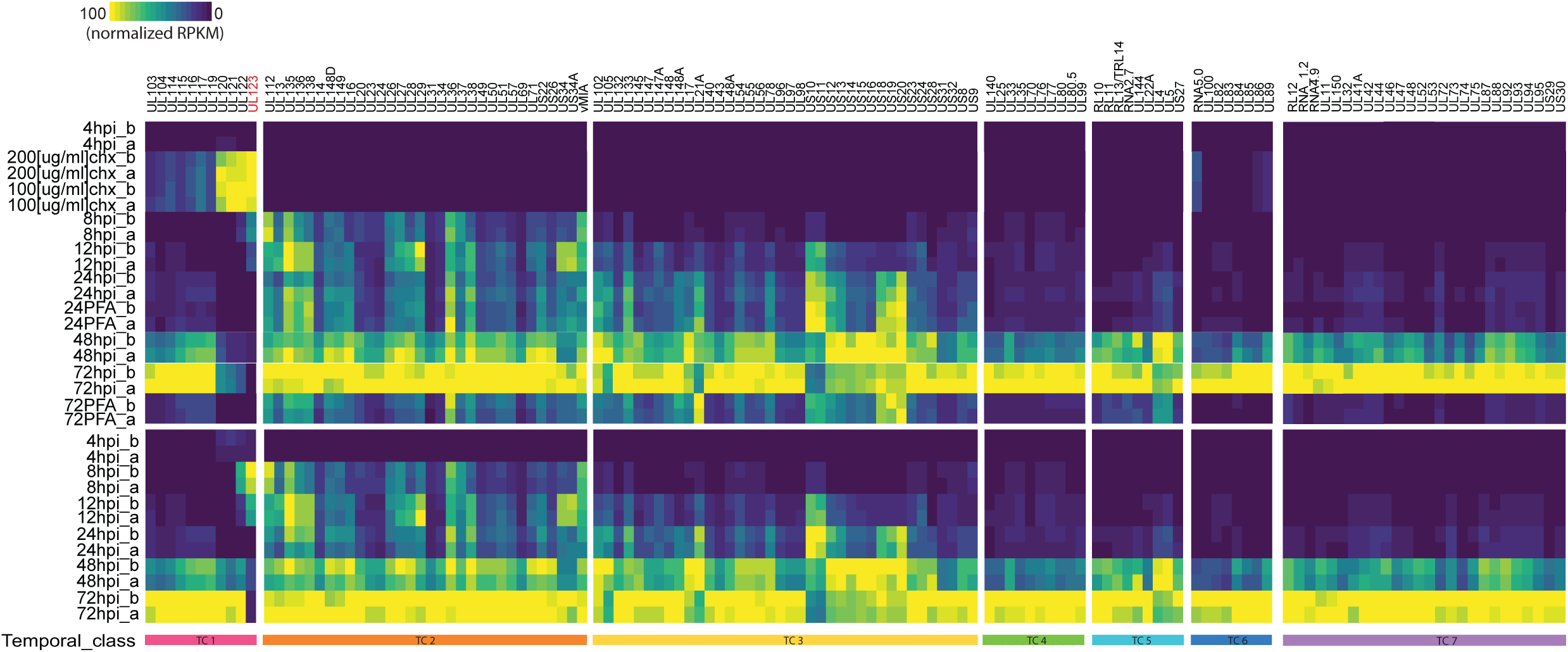
RPKM values of HCMV genes along infection of fibroblasts. Heatmap depicting absolute levels of viral RNA transcripts in reads per kb per million (RPKM) in HCMV infected fibroblasts. Expression patterns of viral genes are shown including CHX and PFA samples in the top panel of the heatmap and without CHX and PFA samples in the lower panel of the heatmap.

**Fig S3.**
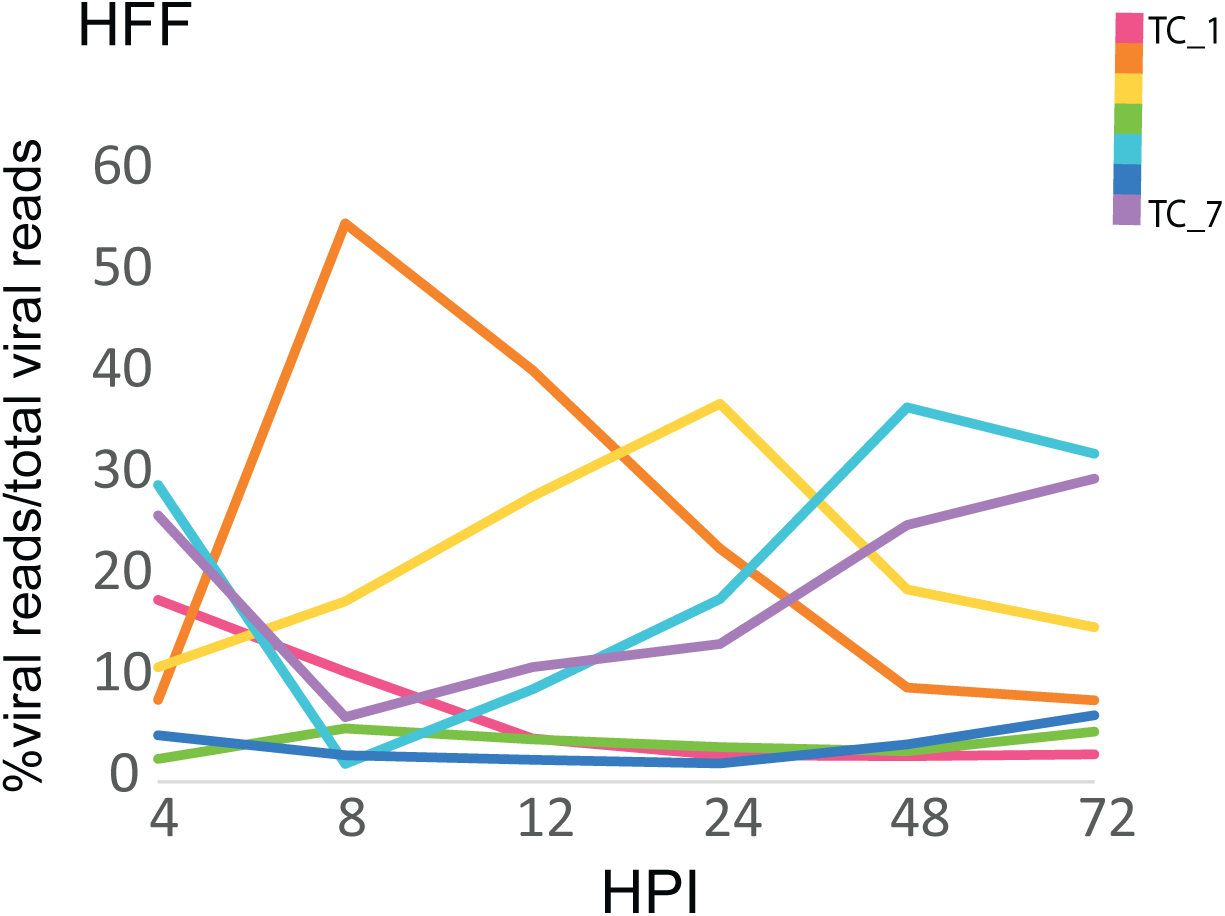
Temporal class dynamics along infection in fibroblasts. Expression Profile of all temporal classes (TCs) along HCMV infection of HFFs, as calculated by percentage of reads from all viral genes in a TC out of all viral reads. Mean values from biological replicates are shown.

**Fig S4.**
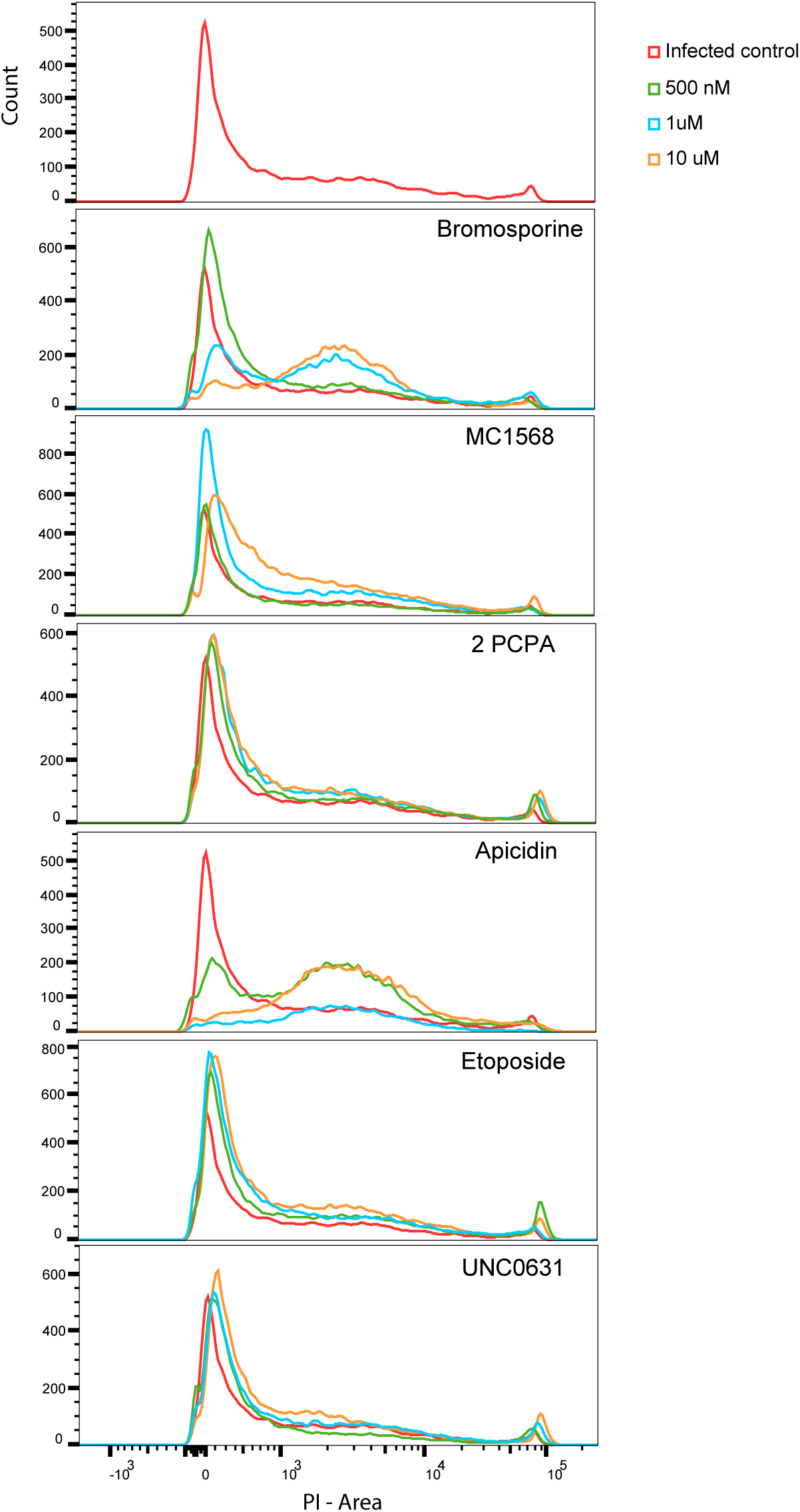
Dose-dependent cytotoxicity of different epigenetic inhibitors. At 24 hours post infection, HCMV infected CD14+ monocytes were incubated with six select drugs from the epigenetic inhibitor library at increasing concentrations (500nM,1uM,10uM). Cell death was measured in an infected control sample and in infected, inhibitor treated samples by propidium iodide staining and flow cytometry.

**Fig S5.**
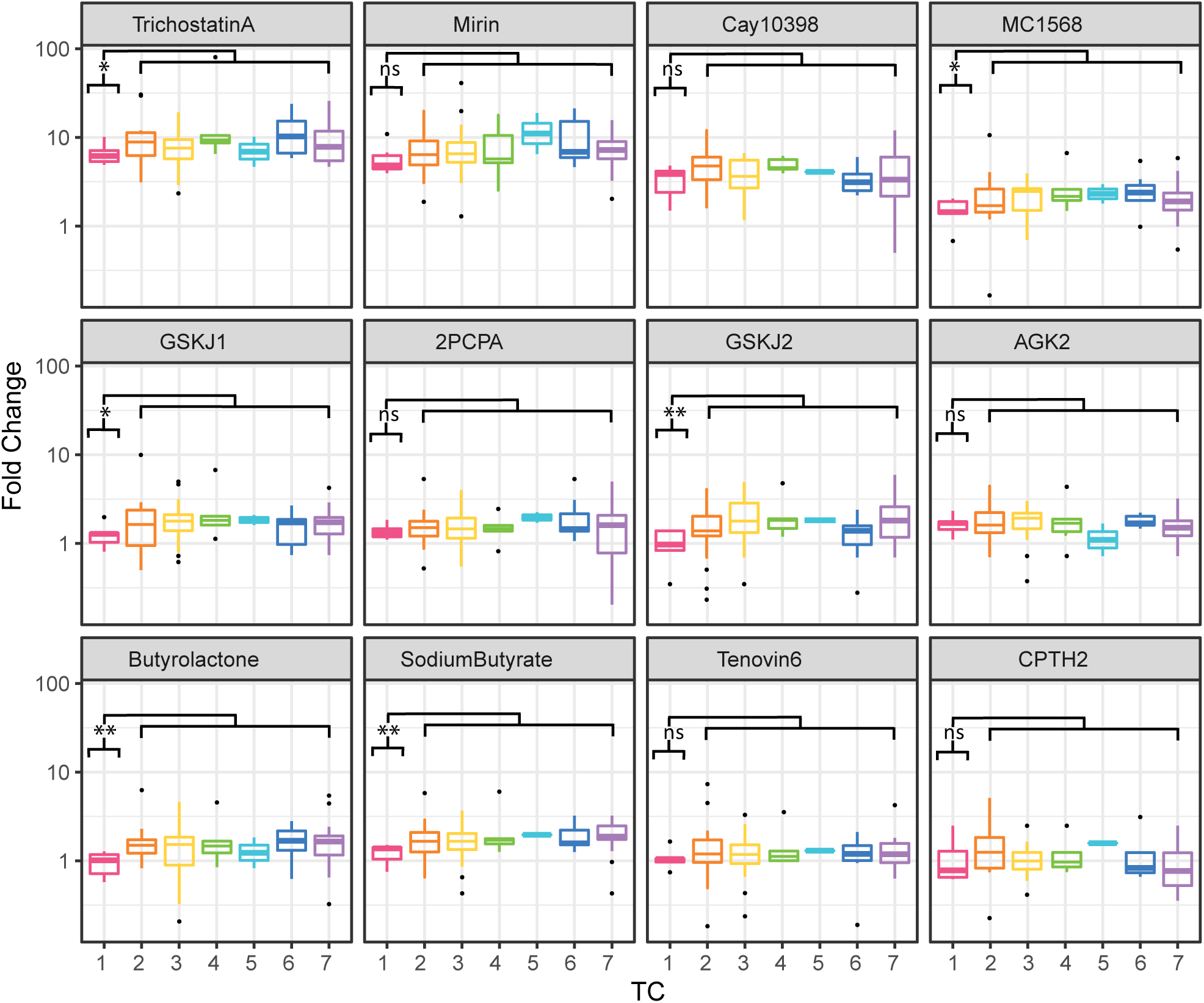
Epigenetic drug treatments reveal unique repression of TC1, immediate early genes. Boxplot showing the transcript level fold change of viral genes from each temporal class between inhibitor treated and control treated HCMV infected CD14+ monocytes. p-value calculated by t-test comparing TC1 and all other TCs is indicated.

**Fig S6.**
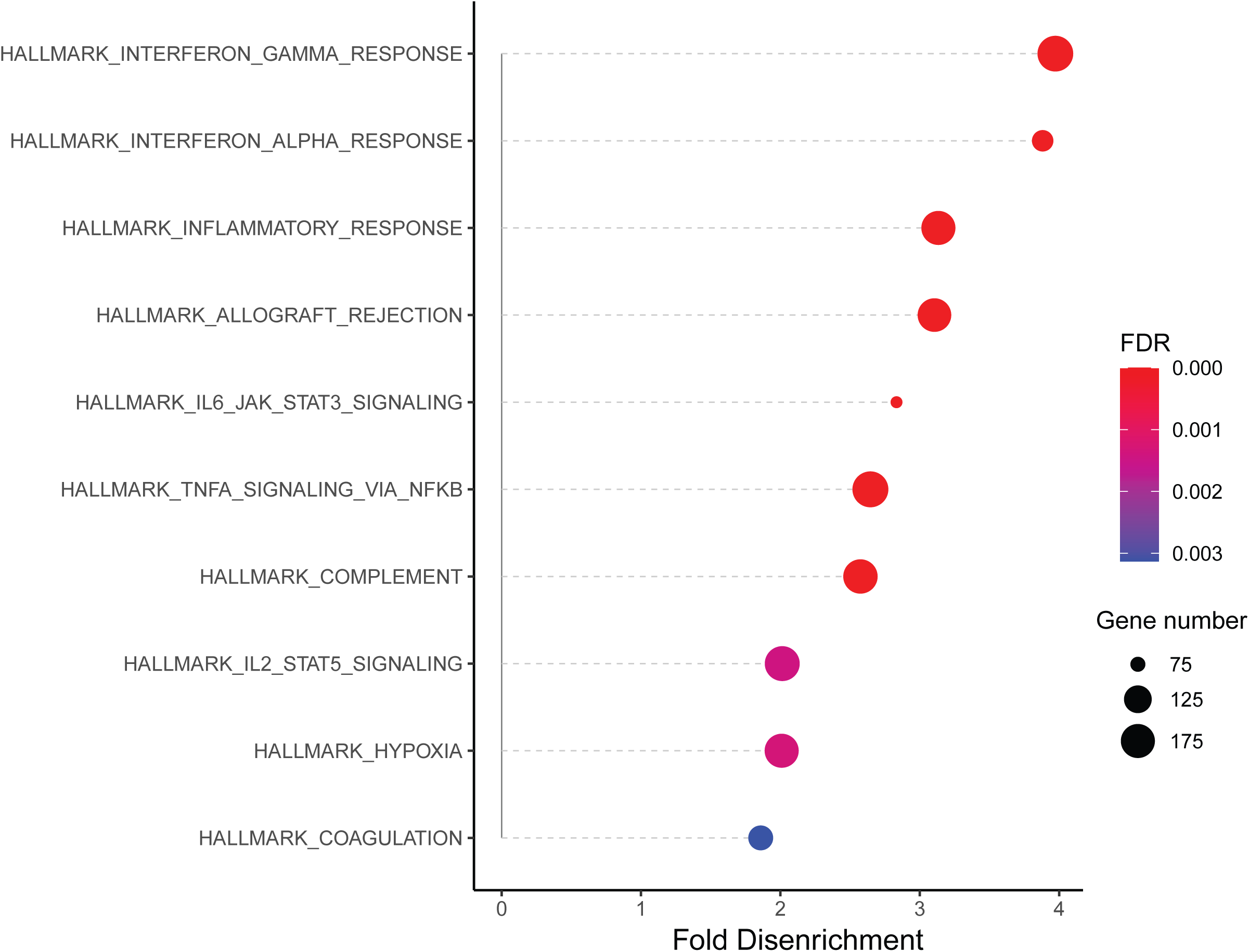
Induction of viral gene expression by TSA time course leads to reduction in expression of immune related genes. Bubble plot showing enriched human hallmark signature pathways which are downregulated following TSA treatment of infected CD14+ monocytes.

